# Gene expression profile dynamics of earthworms exposed to ZnO and ZnO:Mn nanomaterials

**DOI:** 10.1101/2025.06.16.660036

**Authors:** Henk J. van Lingen, Changlin Ke, Edoardo Saccenti, Zoi G. Lada, Marta Baccaro, Nico W. van den Brink, Maria Suarez-Diez

**Affiliations:** Laboratory of Systems & Synthetic Biology, Wageningen University & Research, Wageningen, The Netherlands; Department of Chemistry, Laboratory of Applied Molecular Spectroscopy, Institute of Chemical Engineering Sciences (ICEHT), Foundation for Research and Technology-Hellas (FORTH/ICE-HT), University of Patras, Patras, Greece; Division of Toxicology, Wageningen University & Research, Wageningen, The Netherlands

## Abstract

Zinc oxide containing nanomaterials may elicit toxic responses in environmental organisms such as earthworms. Although toxic responses in earthworms have been reported, few studies have attempted to understand the molecular mode of action dynamics explaining the observed toxicity at a transcriptional level. This study investigates the time-dependent gene expression response in earthworms exposed to ZnO nanomaterials and ZnO:Mn multicomponent nanomaterials. Earthworms were exposed to ZnO or ZnO:Mn for 7 days and to ZnO:Mn or MnCl2 for 14 days. Strong differential gene expression responses were observed after 4 days of exposure to ZnO nanomaterials and after 2 days of exposure to ZnO:Mn. Moderate differential gene expression responses were observed after 14 days of exposure to ZnO:Mn and MnCl2. Gene ontology (GO) enrichment analysis revealed that differentially expressed genes in earthworms exposed to ZnO were associated with terms such as actin, (striated) muscle cells, contractile fiber, myofibril, sarcomere and supramolecular cellular components. In addition, genes that were upregulated after 2 days of exposure to ZnO:Mn were linked to GO terms including cilium, microtubule, cell projection, axoneme and sperm flagellum cellular components. Downregulated genes were enriched to GO terms related to ribosomes, mitochondria, translation, peptide processes, respiration and oxidative phosphorylation. Finally, for earthworms exposed to ZnO:Mn and MnCl2 for 14 days, only a limited number of differentially expressed genes were involved in GO terms related to diverse biological implications. In summary, exposing earthworms to the ZnO and ZnO:Mn nanomaterials elicited a transient response in differential gene expression related to muscle biology and energy metabolism and translation that had largely disappeared by day 14 from the start of the exposure.

## Introduction

Metal oxide nanomaterials (NM) such as those containing zinc oxide (ZnO) have become increasingly common resources for a variety of industrial applications (Papadiamantis *et al*., 2024). Applications of ZnO may include solar cells and semiconductors due to their electrical properties (O zgu r *et al*., 2010), cosmetic products (Lu *et al*., 2015), UV-blockers in sunscreens (Serpone *et al*., 2007) and various biomedical applications such as fluoresence bioimaging and anticancer activities (Prashanth *et al*., 2024). Although ZnO nanomaterials are highly effective, loading them with transition metals such as manganese generates MultiComponent NanoMaterials (MCNMs) with more distinctive electronic properties. This increases the versatility for applications such as biocatalysis (Aulakh *et al*., 2022) and fuel cell technology (Hasan *et al*., 2024). Due to the potential utility of ZnO-containing MCNM, their toxicity to humans and environmental ecosystems has been screened, but a mechanistic understanding has not yet been obtained (Zhang *et al*., 2022).

Environmental impact and toxicity of nanomaterials to the soil ecosystem has previously been assessed using earthworms. Exposure of earthworms to Ag2S nanoparticles at 10 mg Ag kg^−1^ of soil for 28 days did not affect the burrowing behavior at the used exposure content (Baccaro *et al*., 2019). In addition, exposure earthworms to bimetallic nanoparticles and combinations of nano- and ionic forms of Au and Ag at nominal contents of 25 mg Ag kg^−1^ and 1.5 mg Au kg^−1^, indicated that an increased dissolution associated with ionic forms was a driving factor for metal uptake in earthworms (Baccaro *et al*., 2022). Co-exposure to both metals in different forms showed different accumulation patterns compared to the single metal exposure, which points to the importance of toxicity testing of chemical mixtures. Furthermore, exposure of earthworms to ZnO NM and ZnCl2 at contents up to 750 mg Zn kg^−1^ for over 28 days showed ZnO did not have a clearly harmful effect, whereas ZnCl2 was toxic as indicated by significant reduction in survival and the subsequent impact on reproductive efficiency (Singh *et al*., 2024). In line with the latter results, a lower growth constant and lower cellular respiration rate was observed for ZnCl2 than ZnO after exposure for 98 days (Filipiak *et al*., 2021). Nevertheless, ZnO nanoparticles may act as a reproductive toxicant to earthworms at contents ≥ 500 mg kg^−1^, although the exact effect depends on the soil properties, with the toxic reproductive potential of ZnO nanoparticles in clayey soil being greater than in sandy soil (Samarasinghe *et al*., 2023).

Although various studies have reported commonly applied toxicity tests using earthworms, few studies have attempted to understand the molecular mode of action explaining the observed toxicity. This mode of action in earthworms exposed to ZnO- containing NM may be assessed through gene expression quantification approaches. Novo *et al*. (2020) found expression changes of genes encoding for several Zn transporters after 28 day exposure to ZnO NMs and ZnCl2 (*i.e*. Zn in the ionic form) at EC50 values ranging from 200–600 mg Zn/kg soil. Furthermore, specific gene expression associated with ZnO NM exposure was related to vesicular transport and specifically to endocytosis.

Subsequent analysis indicated enhanced adhesion and peptidase activities linked to a normal physiological function of ionic Zn during epithelial–mesenchymal transition. In addition to the latter relatively long-term exposure study, Gomes *et al*. (2022) observed the gene expression response in potworms after 1 to 4 days of exposure to ionic Zn from ZnCl2 at contents of 93 and 145 mg Zn/kg soil. The response was time-dependent with a clear peak in number of differentially expressed genes (DEG) on day 3. These DEG indicated upregulated zinc transporters, oxidative stress and effects on the nervous system. Moreover, few genes were differentially expressed on multiple days, which indicated a cascade of molecular events occurring in time rather than a sustained response.

The relatively small number of studies that assessed the mode of action in earthworms exposed ZnO nanomaterials and the absence of studies exploring the gene expression response in earthworms exposed to ZnO-containing MCNM warrants further investigation. Additionally, the time-dependent gene expression response observed by Gomes *et al*. (2022) suggests that an experimental design in which samples are collected on multiple days from the start of the exposure gives additional insights into molecular responses in exposed earthworms. This study aims to investigate the dynamics of the response in earthworms exposed to ZnO and ZnO:Mn nanomaterials through gene expression analysis.

## Materials & Methods

### Nanomaterials, earthworms, soil jar preparation and experimental design

We present three experiments in which earthworms were exposed to ZnO NM (experiment 1) or to ZnO:Mn MCNM (experiments 2 and 3). Treatments, sampling days and sample sizes of the design of these experiments are schematically represented in Fig. 1. In the first experiment, we exposed earthworms, ordered as *Eisenia fetida* from Dutchworms (Oostwold, The Netherlands), to either control or ZnO nanomaterial (Phornano Holding GmbH; Korneuburg, Austria) treated LUFA 2.2 soil (LUFA Speyer, Germany). The supplier indicated the soil was air-dried and sieved (<2 mm) and had a pH of 5.6 ± 0.29 (mean ± SD), an organic carbon content of 1.82 ± 0.48 w/w%, a measured cation exchange capacity of 9.54 ± 1.36 meq/100 g, and a water holding capacity of 48.9 ± 5.6 g per 100 g. Glass jars were prepared with air dried soil (150 g per earthworm) spiked with ZnO to reach a nominal content of 850 mg/kg and then additional deionized water (20% w/w, ∼47% water holding capacity) was added. Treatment soils were. Preparing control and treatment soils, adding the ZnO nanomaterial and water, and the soil mixing was performed manually. This resulted in 16 jars with control soil and 16 jars with ZnO treated soil. After 4 days of equilibration of the soil, jars were refilled with deionized water up to their weight at soil preparation, after which 4 clitellated earthworms, *i.e*. sexually matured individuals, with an average weight of 386 ± 39 mg were randomly placed into every jar. Additionally, an amount of 0.5 g of horse manure per worm was amended on top of the soil in the jars to feed the earthworms. This horse manure was obtained from an organic farm (Bennekom, The Netherlands) with known absence of pharmaceutical use.

**Figure 1.**
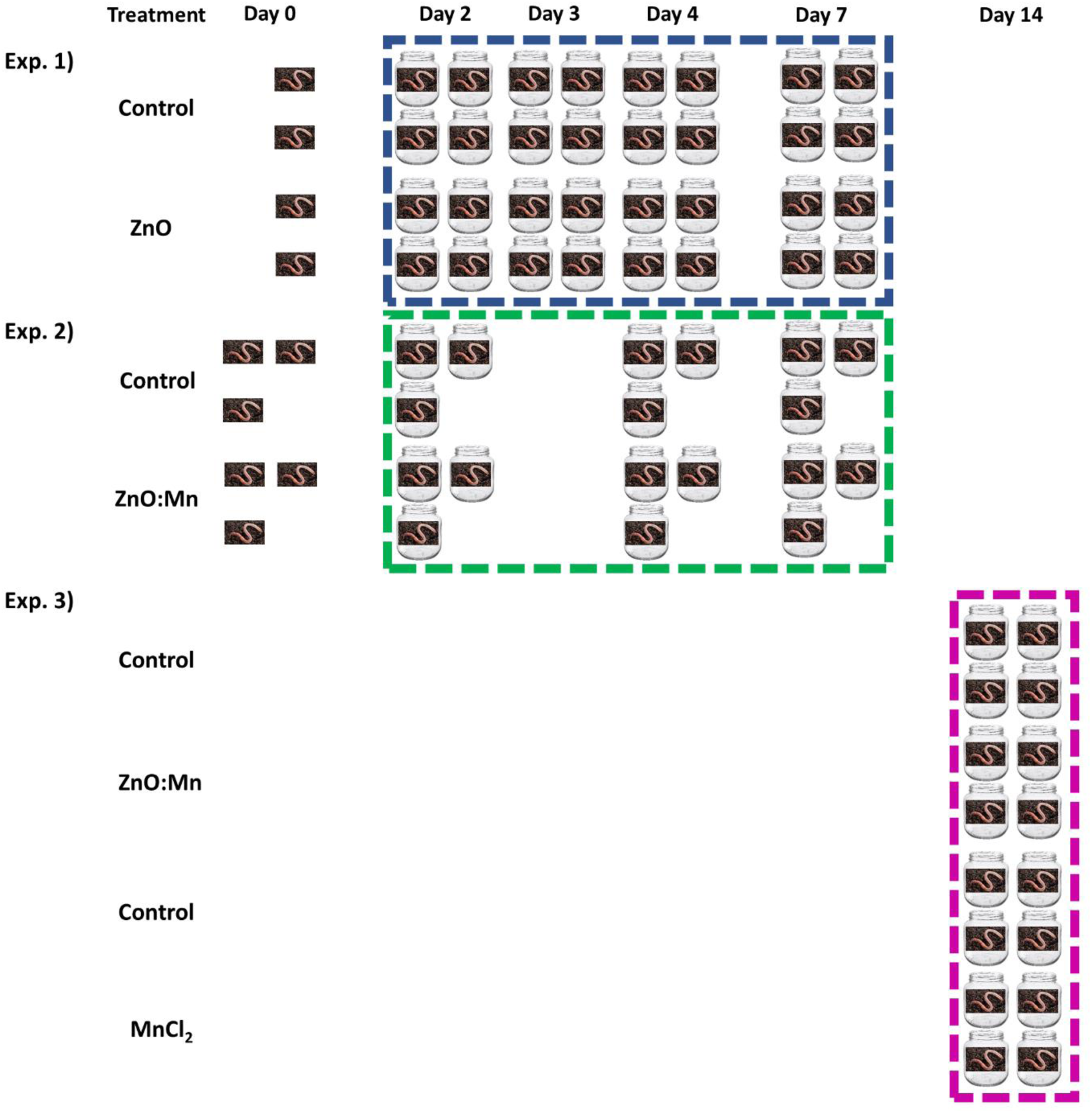
Treatments, sampling days and sample sizes in earthworm experiments 1, 2 and 3. Dotted squares with jars indicate incubated earthworms. Earthworms shown outside the dotted squares were sampled as reference worms before the start of the incubation on day 0 in experiments 1 and 2.

At the start of the first experiment, 4 earthworms were snap frozen in liquid nitrogen and stored at -80 °C. Then, throughout the entire experiment, jars containing the prepared soils and earthworms were stored in a climate-controlled cabinet (Weiss Technik, Germany) at 20 °C, 75% air humidity and a light intensity of 30 µmol/m^2^/s. Lights were switched on for 24 hours per day to stimulate the earthworms to stay in the soil and avoid that they would escape from the jars. All 32 jars stayed in the cabinet for 7 days and were positioned in a completely staggered manner with respect to treatment and time of earthworm sampling using a rectangle of 4 rows and 8 columns. Sampling took place on days 2, 3, 4 and 7 from the start of the exposure. Hence, on every sampling day, earthworms from 4 jars with control soil and 4 jars with ZnO treated soil were collected, carefully rinsed using deionized water, snap frozen in liquid nitrogen and then stored at - 80 °C. In addition, approximately 35 g of soil was collected per jar and then oven dried at 70 °C for 24 hours. Furthermore, throughout the experiment, jars were refilled with deionized water up to the total weight that every jar with soil and its earthworms had at the start of the exposure. This water refill was repeated every incubation day apart from weekend days. After the 7 day exposure period had finished, one collected whole earthworm per jar was taken from the freezer and ground in liquid nitrogen using a mortar and pestle. The ground tissue was then divided into two equal amounts

The second experiment is also represented in Fig. 1 and exposed earthworms, also ordered as *Eisenia fetida* from Dutchworms, to either control or ZnO:Mn multicomponent nanomaterial (Phornano Holding GmbH; Korneuburg, Austria) treated LUFA 2.2 soil. Besides a different nanomaterial and a different number of jars, the experimental conditions were the same as in experiment 1. These other differences applied to the number of jars and earthworms, and the sampling days. In the second experiment, 9 jars with control soil and 9 jars with ZnO:Mn treated soil at 850 mg nanomaterial/kg soil were prepared. Thereafter, 8 clitellated earthworms were snap frozen in liquid nitrogen and then stored at -80 °C and 2 clitellated earthworms with an average weight of 604 ± 63 mg per earthworm were randomly placed into every jar. All 18 jars stayed in the cabinet for 7 days and were positioned in a staggered manner with respect to treatment and time of earthworm sampling using a rectangle of 3 rows and 6 columns. These samplings took place on days 2, 4 and 7 from the start of the exposure. Hence, on every sampling day, earthworms from 3 control jars and 3 ZnO:Mn treated jars were collected.

Experiment 3 was designed with a 14-day exposure period, representing the endpoint measurement relative to Experiments 1 and 2, and is schematically illustrated in Fig. 1. This third experiment of the present study exposed earthworms, also ordered as *Eisenia fetida* from LASEBO (Nijkerk, The Netherlands), to either control or ZnO:Mn multicomponent nanomaterial (Phornano Holding GmbH; Korneuburg, Austria) treated LUFA 2.2 soil, or to either control or MnCl2 treated LUFA 2.2 soil. The nominal exposure content of ZnO:Mn was also set at 850 mg/kg soil, which aimed at quantifiable zinc above the natural background content in the soil. Noting the 95:5 ratio of ZnO:Mn of the MCNM, the nominal exposure content of ionic Mn from MnCl2 was set at 45 mg Mn/kg soil. 4 jars with control soil and 4 jars with ZnO:Mn treated soil, and 4 jars with control soil and 4 jars with MnCl2 treated soil were prepared. 6 clitellated earthworms with an average weight of 584 ± 64 mg were randomly placed into every jar. All 16 jars stayed in the cabinet for 14 days, after which all earthworms were sampled at once, depurated from their gut content for 24 hours on a petri dish, and then snap frozen in liquid nitrogen and stored at -80 °C. One collected whole earthworm per jar was taken from the freezer and ground in liquid nitrogen using a mortar and pestle. The ground tissue was then divided into two equal amounts enabling transcriptomics analysis. Other details regarding experimental design and sampling were as in the first experiment in which earthworms were exposed to ZnO.

### Nanomaterial characterization and Determination of Zn and Mn in soil

Raman spectroscopy of ZnO samples showed the most intense peak at 438 cm^−1^ corresponding to the E₂ mode typical for the hexagonal crystal structure of ZnO. Additional less intense peaks were observed at 330 and 580 cm^-1^, and a broad peak around 1150 cm^-1^. For ZnO:Mn, a less intense 438 cm^-1^ peak in the Raman spectrum indicates defects and a breakdown in translational symmetry caused by substitutional or interstitial Mn dopant. Additional peaks at 530, 570, and 675 cm⁻¹ only observed after Mn doping of ZnO may reflect a local surface distortion that could influence the reactivity of the nanoparticles by (Hu *et al*., 2011). Furthermore, X-ray diffraction analysis confirmed the hexagonal structure of ZnO and showed the doping process reduces the overall crystallinity of the ZnO structure and broader diffraction peaks pointed to a hexagonal wurtzite (*P*63*mc*) lattice symmetry (Safaei-Ghomi and Ghasemzadeh, 2017). The Mn dopant interferes with the lattice during synthesis causing more disordered region and reducing the crystal symmetry and increase the crystallite size. In addition, scanning electron microscopy image analysis evaluating the crystal morphology indicated ZnO appears in particles that had a diameter of 50 ± 19 nm, whereas ZnO:Mn has a surface structure, suggesting increased aggregation or particle fusion due to Mn incorporation. Diameter size of ZnO:Mn could not be determined due to its surface morphology. Further details of the experimental characterization of the nanomaterials are provided in the Supplementary information, along with the Raman and X-ray diffraction spectra and scanning electron microscopy images (Fig. S1-S3). The present observed lattice strain and defects, and morphological changes induced by Mn doping enhance chemical reactivity and bioavailability of the nanoparticles. These properties may influence the extent and mechanisms of biological interaction, including uptake and oxidative stress responses, ultimately inducing cellular toxicity profile (Persaud *et al*., 2020). For further details on the ZnO nanomaterials and ZnO:Mn MCNM, which is the Mn doped analogue of ZnO, we refer to Carreira-Barral et al., (2025) and the Supplementary information.

All 32 and 18 oven dried soil samples from emptied jars in experiments 1 and 2, respectively, were ground at 500 µm and then dissolved using aqua regia (3:1 ratio of HCl:HNO3). Then, after digestion and atomization, the concentrations of the corresponding elements were determined using inductively coupled plasma optical emission spectrometry (Thermo iCAP-6500 DV, Thermo Fisher Scientific; ICP-OES). The ICP-OES measurements were conducted in accordance with the guidelines in NPR-6425 and NEN-6966. Internal standards elements of the ICP-OES analysis included Be, Sc and Y. Grinding, further processing and metal content determination were performed by the soil chemistry laboratory (CBLB) at Wageningen University. Zn and Mn extraction from the 16 soil samples from experiment 3 was performed using Teflon vessels and 7 mL of reverse aqua regia (3:1 ratio of HCl:HNO3; Merck, Darmstadt) on hot plates in an open system. After diluting properly, the digested samples were analyzed by inductively coupled plasma-mass spectrometry (ICP-MS Nexion 350D, Perkin-Elmer Inc., Waltham, MA). The calibration curve was prepared from solutions of Zn^2+^ and Mn^2+^ (standard stock solution 1000 mg/L of ionic metal ion, Merck, Darmstadt) in a matching matrix.

### RNA extraction and sequencing

Ground earthworm samples from experiments 1 and 2 were sent to BMKGene (Mu nster, Germany) for RNA extraction and sequencing. A TRIzol kit was used to complete the extraction of RNA, after which the concentration of extracted nucleic acid was detected using a Nanodrop2000 spectrophotometer (Thermo) and the integrity with an Agient2100 Bioanalyzer LabChip GX (Perkin Elmer). Libraries from poly(A) enriched RNA were prepared along purification, fragmentation, reverse transcription to cDNA, ligation and amplification using a Hieff NGS Ultima Dual-mode mRNA Library Prep Kit for Illumina (Yeasen; model:13533ES96) OR VAHTS Universal V8 RNA-seq library preparation kit for Illumina – NR605. Prepared libraries were inspected by Qsep-400 and then sequenced on an Illumina Novaseq X platform (Illumina, San Diego, CA) and reads were generated using paired-end 2×150 base pairs.

Ground earthworm samples from experiment 3 were sent to GENEWIZ Germany GmbH (Leipzig, Germany) for RNA extraction and sequencing using the same requirements as for the orders of experiments 1 and 2 sent to BMKgene. Qiagen RNeasy Mini kit (Qiagen, Hilden, Germany) was used to complete the extraction of RNA. The concentration of extracted RNA was detected using a Qubit 4.0 Fluorometer (Life Technologies, Carlsbad, CA, USA) and the integrity was checked with RNA Kit on Agilent 5300 Fragment Analyzer (Agilent Technologies, Palo Alto, CA, USA). Libraries from poly(A) enriched RNA were prepared along purification, fragmentation, reverse transcription to cDNA, ligation and amplification using the NEBNext Ultra II RNA Library Prep Kit for Illumina (NEB, Ipswich, MA, USA). Sequencing libraries were validated using NGS Kit on the Agilent 5300 Fragment Analyzer (Agilent Technologies, Palo Alto, CA, USA), and quantified using a Qubit 4.0 Fluorometer (Invitrogen, Carlsbad, CA). The sequencing libraries were multiplexed and loaded on the flow cell on the Illumina NovaSeq 6000 instrument using paired-end 2×150 base pairs.

### Raw reads quality control, read alignment and quantification for all samples

The raw Illumina paired-end reads were processed using the workflow provided in the Supplementary information and at: https://workflowhub.eu/workflows/336?version=1, which was developed with the Common Workflow Language (CWL) v1.2. The workflow was used with default settings plus rRNA removal based on ribo-Kmers from SILVA (Bushnell, 2014; ribo-Kmers were downloaded from: https://portal.nersc.gov/dna/microbial/assembly/bushnell/) to perform quality control, trimming, filtering, and classification to ensure high-quality input for downstream analyses. Optional steps in the workflow were omitted. Assembled and annotated genomes of *Eisenia andrei* and *Eisenia fetida*, which are morphologically similar earthworms, were retrieved from National Genomics Data Center (Genome Warehouse, GWH; http://bigd.big.ac.cn/gwh/) with accession number GWHACBE00000000 (Shao *et al*., 2020) and from NCBI with accession number GCA_003999395.1 (Bhambri *et al*., 2018). Cleaned reads were mapped to the genomes of these species. Samples with mapping rates above 40% were selected for reference-based transcript quantification using FeatureCounts v2.0.1 (Liao *et al*., 2014).

### Reference-free assembly and read counting

Samples with mapping rates to references < 40% were found in experiment 3. For those samples, *de novo* transcriptome assembly was performed using Trinity 2.15.1 (Grabherr *et al*., 2011). Parameters were set to handle strand-specific paired-end data, ensuring accurate transcript orientation. Low-expressed (minimum TPM < 3) and short transcripts (length < 250 bp) were filtered to enhance assembly quality and reduce redundancy. Post-assembly clustering was applied using CD-HIT-EST 4.8.1 (identify threshold = 95) to collapse similar transcripts while preserving biologically relevant isoforms (Li & Godzik, 2006). Coding regions in the output assembly were predicted by TransDecoder 5.5.0 on Galaxy (https://github.com/TransDecoder/TransDecoder). The cleaned reads were then aligned back to the filtered assembly and Trinity-grouped genes and transcripts abundance estimation were conducted with Salmon v0.14.1 (Patro *et al*., 2017). The assembled transcripts were annotated using Trinotate 4.0.2 (Griffith *et al*., 2015) to assign putative functions. The annotation pipeline included diamond 2.1.10 BLAST (Buchfink *et al*., 2015) searches against the SwissProt database (both BLASTX and BLASTP) and Pfam domain identification for protein family assignments. Additionally, EggnogMapper v2.1.12 (Cantalapiedra *et al*., 2021) was used for orthology-based annotations to assign gene ontology (GO) enrichment terms and KEGG pathways.

### RNA-seq data downstream analysis

Prior to statistical testing, genes with low expression levels were filtered out. Low expression was defined as less than 50 reads per gene across samples, except for the 5 samples mapped to *E. andrei* in experiment 3 for which a cutoff of 25 reads per gene across samples was applied. Normalization of the read counts was performed using the relative log expression method using the DESeq2 package v1.30.0 (Love *et al*., 2014) in R version 4.4.2. Considering Day × Treatment interaction effects for experiments 1 and 2, normalized read counts were fitted using the negative binomial model approach implemented in DESeq2. For experiment 3, only the treatment effects were considered for the negative binomial model as all samplings were taken on the same experimental day. Thereafter, Wald tests were conducted to determine the significance of differential expression for each gene. To correct for multiple comparisons, the Benjamini-Hochberg procedure was applied, resulting in adjusted P-values. Genes with a log2 fold change greater than 0.58 or less than -0.58, corresponding to a fold change of 1.5 times or 1/1.5 times and an adjusted P-value less than 0.05 were considered significantly differentially expressed.

Furthermore, a regression-based analysis was performed that used a generalized linear model specifically suited for the estimation of parameters of serial data using time as an independent quantitative (rather than qualitative as used in the differential expression analysis above) variable. The time series data generated for the nanomaterial and control treatments in experiments 1 and 2 were also fitted to a negative-binomial model in this regression-based analysis. The model fitting procedure was performed along a step in which genes were selected, a step that included backward selection of time and treatment variables with non-zero regression coefficients, and a step that only retained Genes of models with R^2^ > 0.7. Following these three steps, retained genes were subdivided into two groups using the hierarchical clustering method. The regression-based analysis and clustering were performing using the maSigPro package version 1.70.0 in R (Conesa *et al*., 2006; Nueda *et al*., 2014).

Functional GO enrichment of genes found differentially expressed or selected using regression-based analysis was performed by the hypergeometric function to model the background probability and the Benjamini–Hochberg procedure was used to correct for multiple testing and generate adjusted P-values. Annotated GO terms were downloaded from the KEGG/Gene Ontology database (Kanehisha, 2016). GO enrichment analyses were performed using the clusterProfiler package in R (Wu *et al*., 2021). R code used for all downstream analysis steps is publicly available at https://git.wur.nl/ssb/publications/gene-expression-profile-dynamics-and-metabolome-of-earthworms-exposed-to-nanomaterials/-/tree/main/Earthworms

## Results and Discussion

As schematically indicated in Fig. 1, three experiments were performed. In experiment 1 earthworms were exposed to ZnO and samples were taken on days 0, 2, 3, 4 and 7 from the start of the exposure. In experiment 2, earthworms were exposed to ZnO:Mn and samples were taken at 0, 2, 4 and 7 days. Finally, experiment 3 represents an endpoint measurement in which earthworms were exposed to ZnO:Mn and MnCl2 for 14 days until samples were taken. The soil Zn contents in experiments 1, 2 and 3 along with the soil Mn content in experiments 2 and 3 were not significantly different from the set nominal contents equal to 650 and 45 mg/kg soil for Zn and Mn, respectively (Fig. S4-S6). In addition, earthworms exposed to ZnO:Mn for 14 days in experiment 3 had Zn contents that exceeded twice the baseline values, whereas Mn content in earthworms exposed to MnCl2 for 14 days showed a more moderate increase (Van Lingen *et al*., 2025). The present study is unique by combining 3 experiments that, individually or taken together, track the transient transcriptional response in earthworms exposed to ZnO nanomaterial and ZnO:Mn multicomponent material along multiple days for a week. Both by treatment and sampling days, these two experiments complement a previous study that quantified gene expression in potworms exposed to ionic zinc in the form ZnCl2 on days 1, 2, 3 and 4 from the start of the exposure (Gomes *et al*., 2022). Furthermore, in addition to studies in which earthworms were exposed to ionic zinc and ZnO nanomaterials for 28 days (Novo *et al*., 2015, 2020), the endpoint measurements after 14 days in our third experiment are complementary by treatment using the ZnO:Mn multicomponent nanomaterial and ionic Mn in the form of MnCl2. The latter MnCl2 and ZnO treatments may be considered positive controls of the ZnO:Mn multicomponent treatment.

### Earthworm transcripts mapped to reference genomes and *de novo* assembly

After considering that specimens of *E. andrei* may be morphologically identified as *E. fetida* (Ro mbke *et al*., 2016), the obtained RNA-seq reads were mapped to both *E. fetida* and *E. andrei* reference genomes. Mapping rates of reads and further bioinformatic metrics per sample for the 3 experiments are available in File S1. Given the mapping rates, samples that mapped > 40% to *E. andrei* and higher to *E. andrei* than to *E. fetida* in experiments 1 and 2 were retained for further downstream analysis (Fig. 2a-b). For experiment 1, this resulted in only 1 control sample being retained for the samples obtained on day 4 from the start of the exposure. Therefore, all control samples obtained for day 4 were excluded. For experiment 3, low mapping rates to both *E. andrei* and *E. fetida* for 11 out of the 16 samples (Fig. 2c) suggested the collection of earthworms included at least three species. In the past, genotyping was performed using cytochrome amplicon sequencing (*e.g*. Huang *et al*., 2007). Here, we present a *de novo* assembly, which is a straightforward alternative that allows capturing the full genetic potential of the (possibly mixed) population. Moreover, it allows developing custom-made annotation files for enrichment analysis. The performed *de novo* assembly resulted in a mapping rate of 94.5% (range from 92.8% to 95.5%) to the constructed reference transcriptome. Correlation heatmaps did not indicate any patterns or suspicion for the retained samples in experiments 1 and 2 (Fig. S7-S8). However, when evaluating the *de novo* assembled reads for experiment 3, the five samples, which mapped the highest to the two *Eisenia* species, correlated rather well to each other, but showed low correlations relative to the remaining eleven samples, which correlated highly to each other (Fig. S9). Hence, a second read count matrix was generated for these five samples based on the *E. andrei* reference genome instead of the *de novo* assembly.

**Figure 2.**
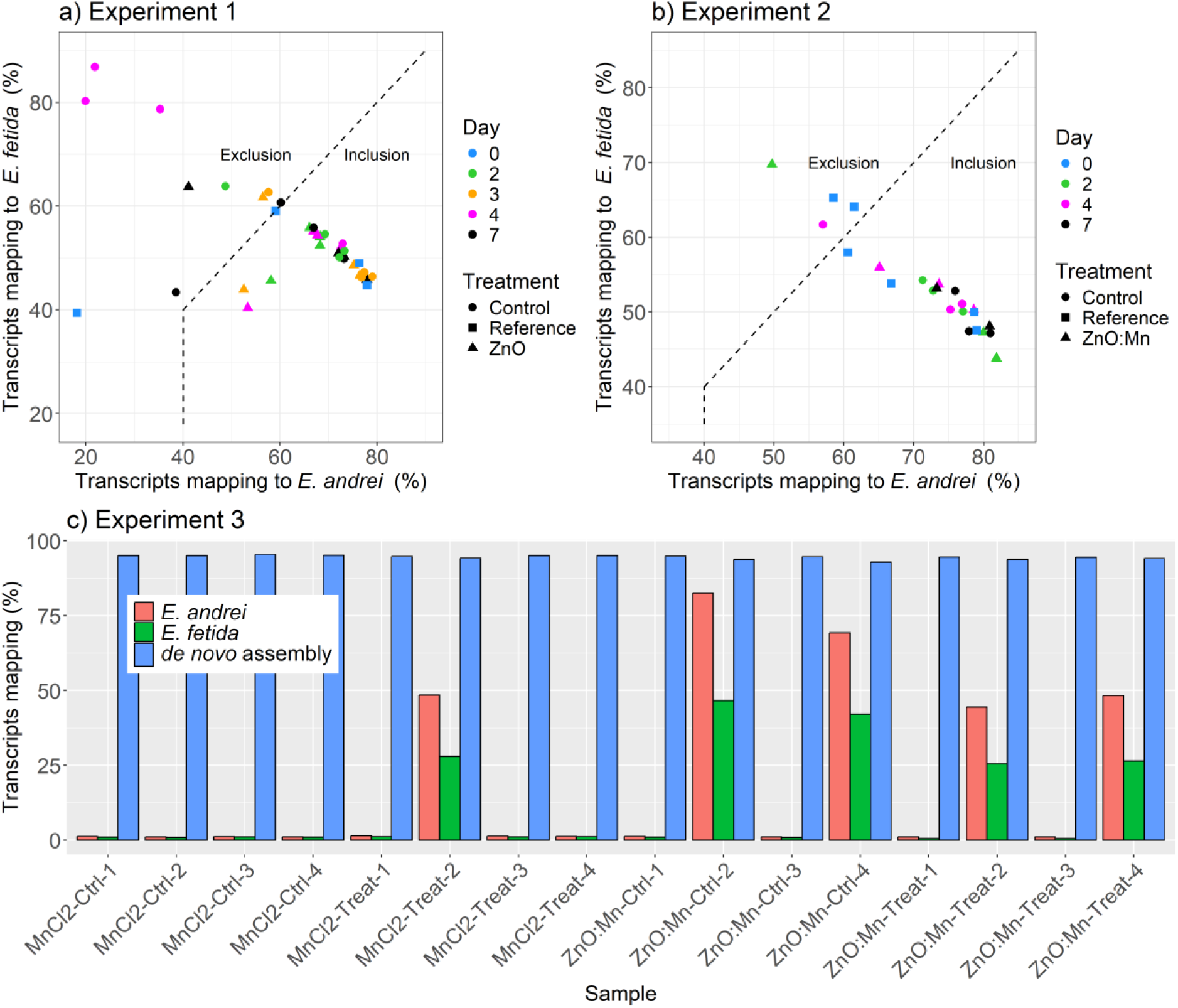
RNA transcript mapping rate per sample to a) *E. andrei* and *E. fetida* in experiment 1, b) *E. andrei* and *E. fetida* in experiment 2, and c) to *E. andrei* and *E. fetida* and a *de novo* constructed assembly in experiment 3. Dashed lines indicate the inclusion and exclusion regions.

### Differential gene expression analysis

DEG were defined by adjusted P-values less than 0.05 and a threshold fold change greater or less than 1.5 and 1/1.5 (corresponding to 0.585 in log2) for identifying biologically meaningful DEG (Schurch *et al*., 2016). The number of DEG in earthworms after 2, 3 and 7 days of exposure to ZnO was very limited (Fig. 3a-c). Out of a total of 28,365 genes, the highest number of DEG observed on day 7 was only 61, of which 8 downregulated and 53 upregulated (Please see File S2 for all DEG). Given that no control samples taken on day 4 were retained this analysis could not be performed on samples from earthworms exposed to ZnO for 4 days. Instead, samples taken at the different days after exposure were compared to the reference samples taken on Day 0. This comparison showed 63, 120, 1037 and 52 DEG along days 2, 3, 4 and 7, respectively, indicating a sharp peak in gene expression on day 4 of which the majority was downregulated (Fig. 4). Furthermore, in line with the latter observation, comparing the ZnO treated samples taken on day 4 to ZnO samples taken on days 2, 3 and 7 also indicated 705, 200 and 282 DEG, respectively (Fig. S10) with the large majority being downregulated. These genes, which were even indicated differentially expressed relative to earthworms samples treated with ZnO taken on days 2, 3 and 7, further indicate a strong temporary gene expression response after 4 days of exposure to ZnO. Although, some caution needs to be taken given the absence of control samples for day 4, a temporary substantial dysregulation of gene expression declining later in time aligns with peak transcriptional responses after 3 days in enchytraeid (potworms) exposed to ZnCl2 (Gomes *et al*., 2022) and earthworms exposed to silver nanomaterials (Tsyusko *et al*., 2012) was observed previously.

**Figure 3.**
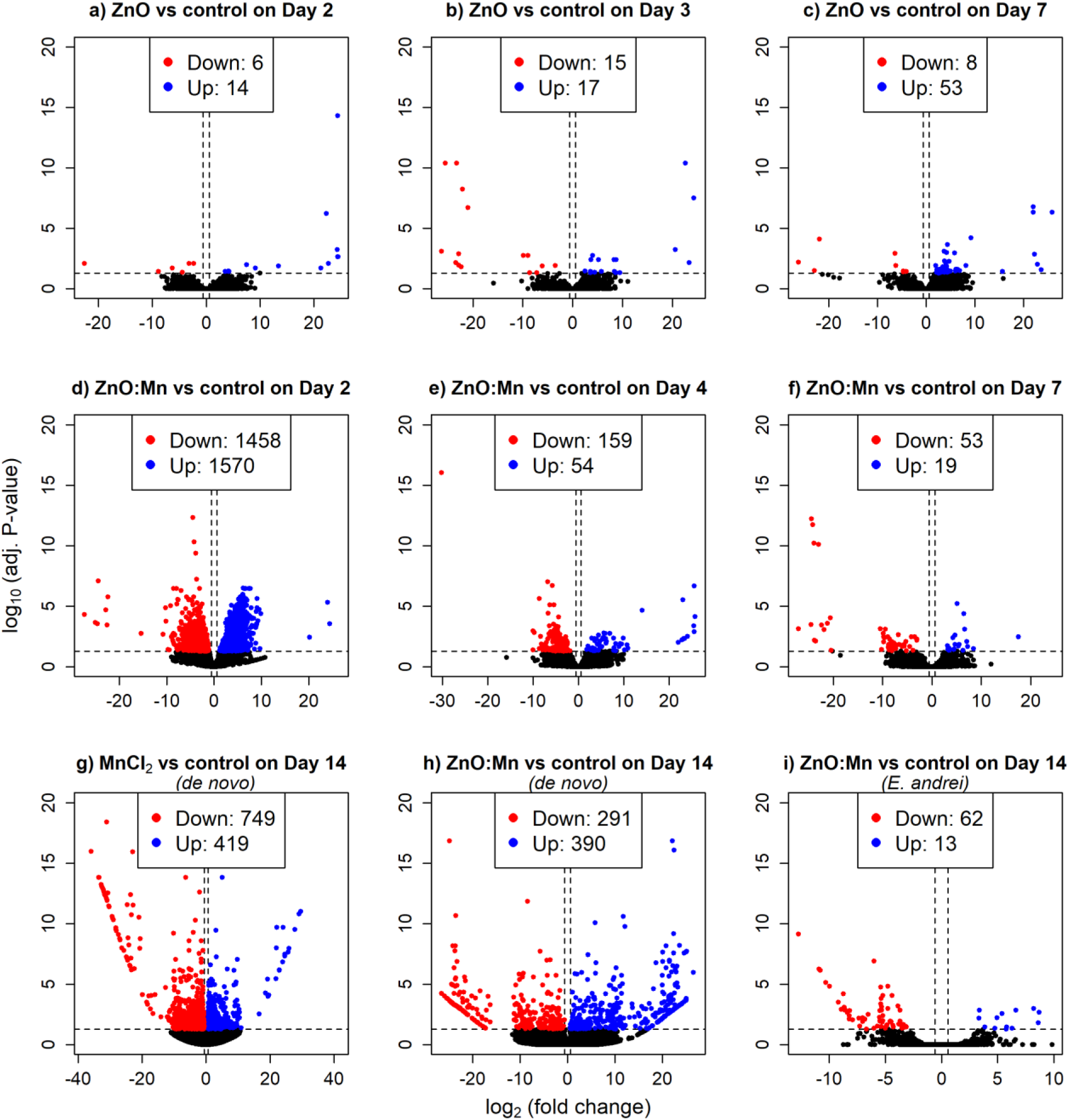
Volcano plots that represent differential gene expression of transcripts mapped to the *E. andrei* reference genome obtained from earthworms exposed to control *vs*. ZnO treated soil for 2, 3 and 7 days in experiment 1 (a-c), from earthworms exposed to control *vs*. ZnO:Mn treated soil for 2, 4 and 7 days in experiment 2 (d-f), of transcripts assembled *de novo* from earthworms exposed to control *vs*. MnCl2 treated soil and control *vs*. ZnO:Mn treated for 14 days in experiment 3 (g-h), and of transcripts mapped to the *E. andrei* reference genome obtained from earthworms exposed to control *vs*. ZnO:Mn treated soil for 14 days in experiment 3 (i).

**Figure 4.**
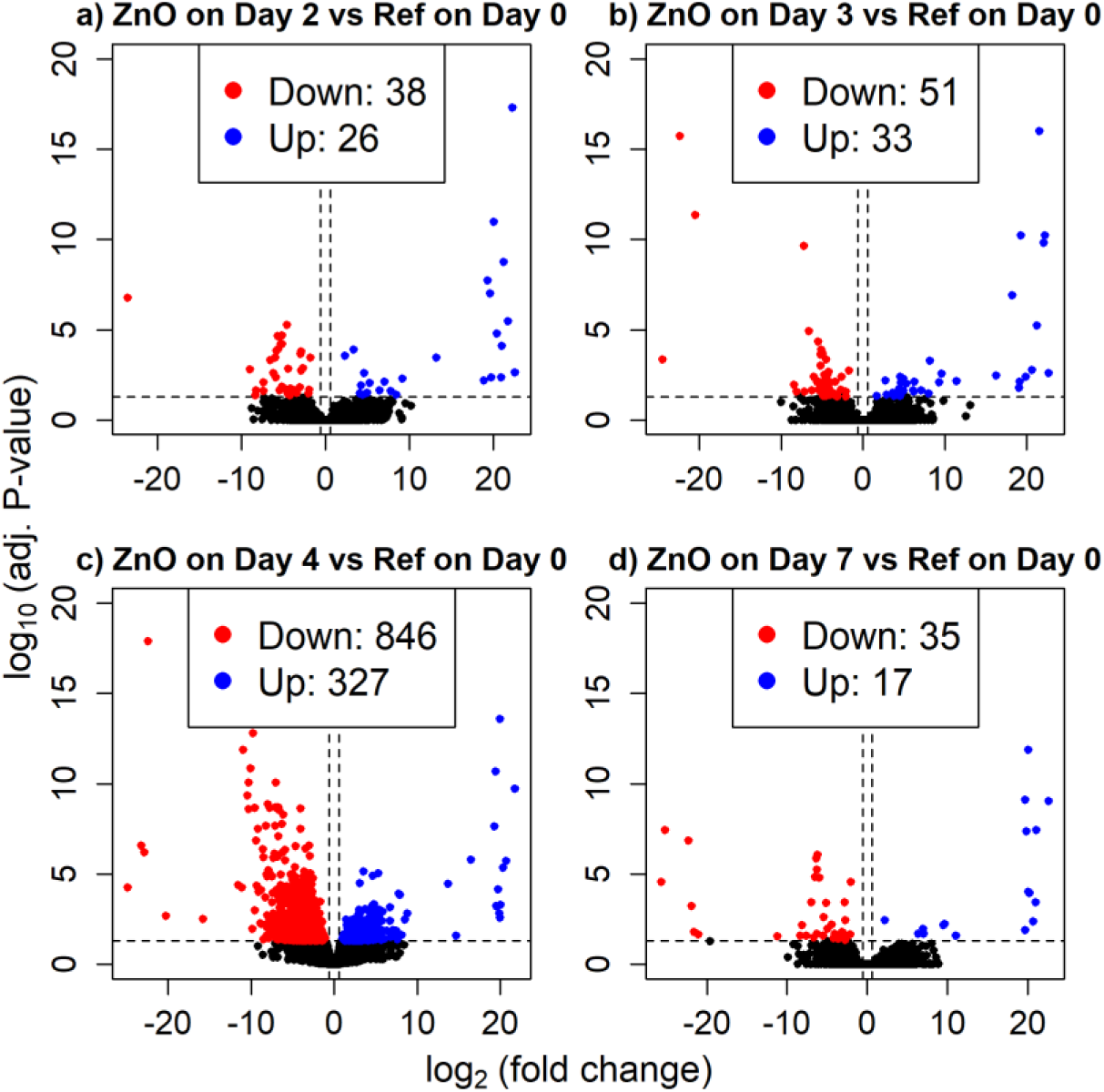
Volcano plots that represent differential gene expression of transcripts mapped to the *E. andrei* reference genome for earthworms exposed to ZnO treated soil for 2, 3 4 and 7 days (a-d) *vs*. earthworms collected before the start of the exposure on day 0 in experiment 1.

Earthworms exposed to ZnO:Mn had more than 3,000 DEG out of 28,241 after 2 days, of which 1458 downregulated and 1570 upregulated (Fig. 3d). This transcriptional response did not persist as only 213 and 72 DEG were observed on days 4 and 7 from the start of the exposure, respectively (Fig. 3e-f). This temporary increase in DEG after 4 days of exposure to ZnO and 2 days of exposure to ZnO:Mn may align with other previous temporary increases in differentially expressed genes such as observed after 3 days of potworm exposure to ionic zinc (Gomes *et al*., 2022) and of earthworm exposure to silver nanomaterials (Tsyusko *et al*., 2012). Exposing earthworms, which are difficult to identify phenotypically, to MnCl2 or ZnO:Mn for 14 days indicated 1168 (749 downregulated, 419 upregulated) and 681 (291 downregulated, 390 upregulated) out of the 46,834 genes annotated in the *de novo* assembled transcriptome, respectively (Fig. 3g-h). These DEG were identified using the eleven samples in experiment 3 showing low mapping to the two *Eisenia* species and were assembled *de novo*. Although Trinity *de novo* assembly may be a powerful approach, which was demonstrated previously (*e.g*. on earthworms exposed to copper; Yu *et al*., 2022), these DEG may be incomparable to DEG of *E. andrei* in experiments 1 and 2. However, reads for the two control and two ZnO:Mn treated samples of experiment 3 mapped to the *E. andrei* reference genome are comparable to gene expression profiles in experiments 1 and 2 by reference genome. These 4 samples indicated only 75 DEG out of 25,984 genes (62 downregulated, 13 upregulated; Fig. 3i).

### Gene expression time-dependent analysis

A regression based approach was used to identify differences in the temporal patterns of gene expression across treatments for experiments 1 and 2. A regression-based generalized linear model was established that evaluated time as a quantitative variable in contrast to the differential expression analysis in which time was considered a qualitative variable. 73 out of 28,365 genes were identified with significant differences in their temporal patters for experiment 1. Clustering resulted in 41 genes that showed increased expression on days 3 and 7 for ZnO treated earthworms relative to the control treated earthworms (Fig. 5, cluster 1A). A second cluster of 32 genes indicated an increase in gene expression in ZnO treated relative to control treated earthworms on days 3 and 7 from the start of the exposure (Fig. 5, cluster 1B). Furthermore, more than 1000 out of 28,241 genes were identified as showing differences in their gene expression temporal dynamics upon exposure to control or ZnO:Mn treated soil in experiment 2. Clustering resulted in a first cluster consisting of 501 genes (Fig. 5, cluster 2A) that contained genes showing increased expression for the ZnO:Mn treated earthworms on day 2 from the start of the exposure, whereas similar gene expression was observed on days 4 and 7. A second cluster of 532 genes indicated downregulated expression in earthworms exposed to ZnO:Mn relative to earthworms that had received the control treatment for 2 days, but this downregulation decreased along days 4 and 7 (Fig. 5, cluster 2B). This difference observed in gene expression for day 2 using regression aligns with the strong transcriptional response indicated by the differential expression analysis on day 2 samples.

**Figure 5.**
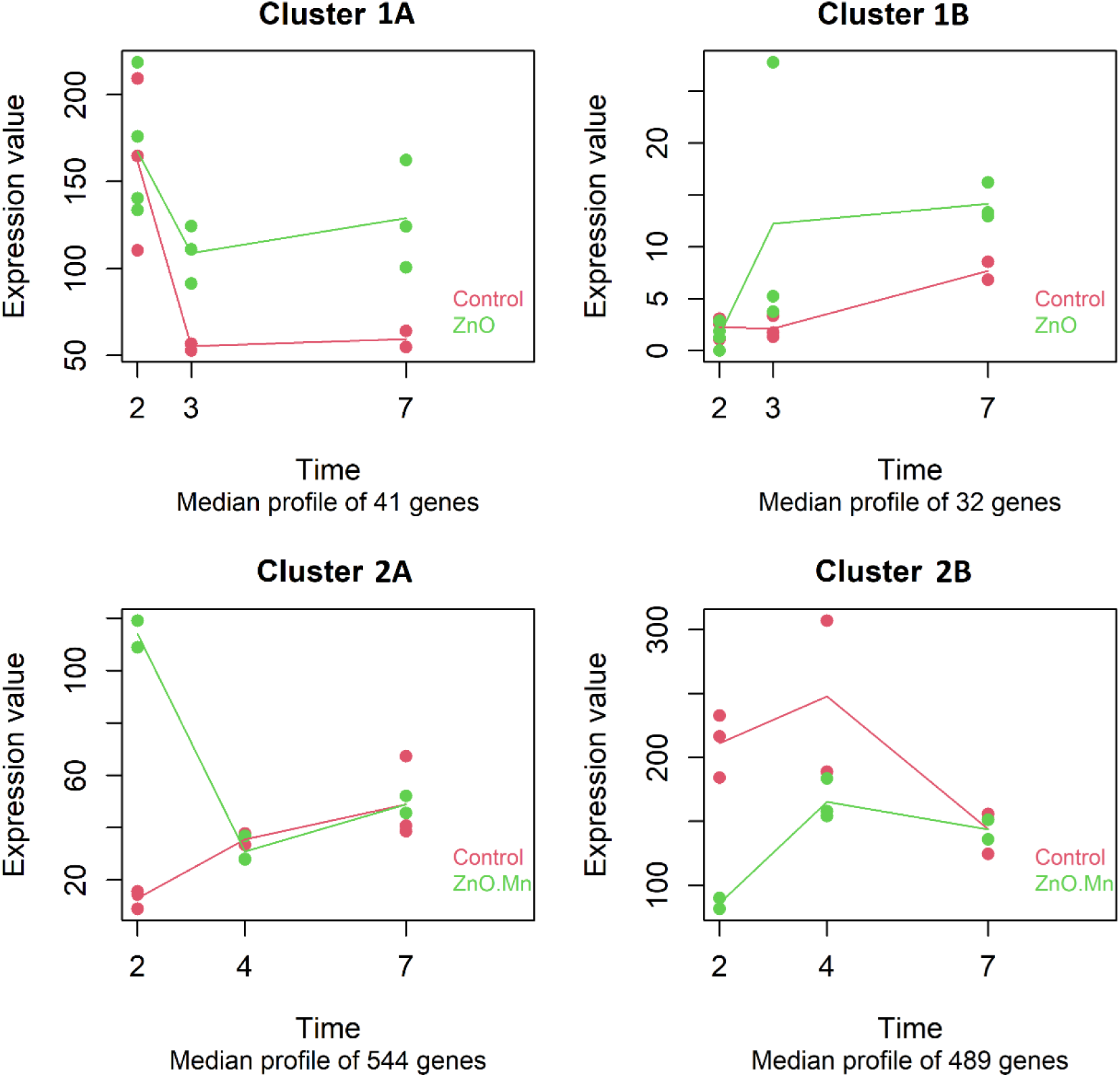
Regression-based profiles of median expression for clusters of selected genes obtained from transcripts mapped to the *E. andrei* reference genome from earthworms exposed to ZnO *vs* control soil for 2, 3 and 7 days in experiment 1 (clusters 1A-B), and exposed to ZnO:Mn *vs* control soil for 2, 4 and 7 days in experiment 2 (clusters 2A-B).

### Gene ontology enrichment

DEG for the various treatments and sampling days underwent enrichment of terms from the Gene Ontology (GO) database (Gene Ontology Consortium, 2019). For earthworms exposed to ZnO, no significant GO-terms were obtained among the DEG at days 2, 3 and 7 from the start of the exposure. For genes downregulated after a 4 days exposure to ZnO, 69 GO terms were found enriched, (adjusted P-values < 0.05) of which the top 15 on adjusted P-value are shown in Fig. 6. GO terms comprised various actin, (striated) muscle cells, contractile fiber, myofibril, sarcomere and supramolecular cellular components. Additional details of the enriched GO terms annotated to the DEG after 4 days of exposure to ZnO are provided in File S3.

**Figure 6.**
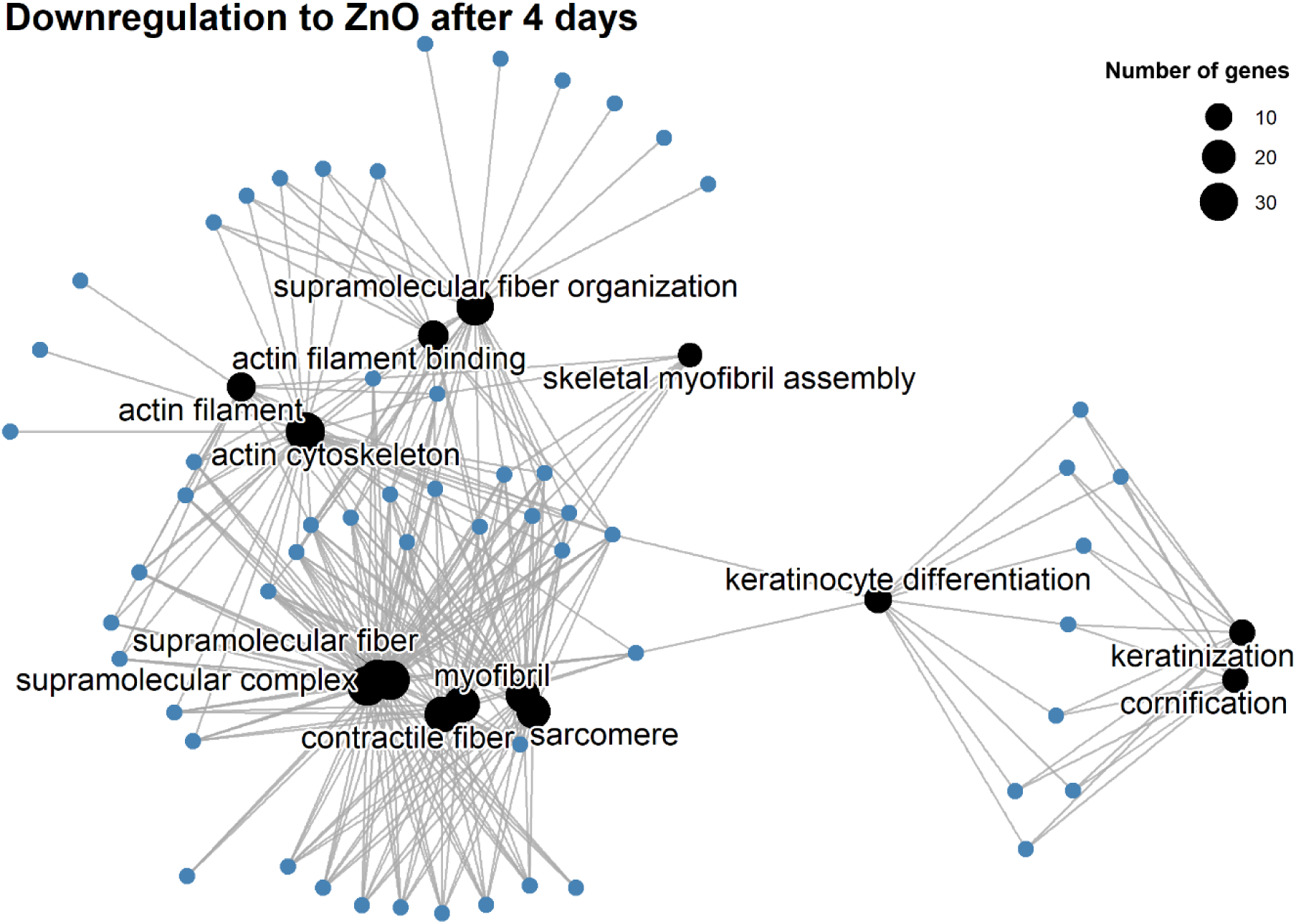
Networks of enriched genes (blue circles) and the top 15 gene ontology terms (black circles, size represents the number of genes associated to each enriched GO term) for downregulated genes in earthworms exposed to ZnO for 4 days in experiment 1. Grey edges connect genes to GO terms.

When considering DEG after 2 days of exposure of earthworms to ZnO:Mn, 54 GO terms were enriched among the upregulated genes and 124 were enriched among the downregulated genes, of which the top 15 are provided (Fig. 7a-b). These 54 and 124 GO terms were associated with up to 61 and 71 genes, respectively. GO terms enriched among the upregulated genes comprised terms associated to various cilium, microtubule-based, cell projection, axoneme and sperm flagellum from the Cellular Component (CC) ontology. In contrast, GO terms enriched among the downregulated genes on day 2 comprised terms from the Biological Process (BP) and Molecular Function ontologies, besides terms from the CC ontology. These GO terms were related to ribosomes, mitochondria, translation, peptide processes, respiration and oxidative phosphorylation. Novo *et al*. (2020) also analyzed transcriptional responses of *E. fetida* earthworms exposed to ZnO NM, but they reported enrichment of GO terms that included vesicular/membrane transport, compound binding, adhesion, extracellular and peptidases, which largely do not align with the present GO terms obtained after 4 and 7 days of exposure to ZnO or 2 days of exposure to ZnO:Mn. Moreover, upregulated genes after 4 days and downregulated genes after 7 days of exposure to ZnO:Mn, showed enrichment of 51 and 96 GO terms, respectively. However, these were associated to no more than 2 genes per GO term and their relevance cannot be ascertained. Additional details of all GO terms enriched among the DEG after 2 days of exposure to ZnO:Mn are provided in Files S3.

**Figure 7.**
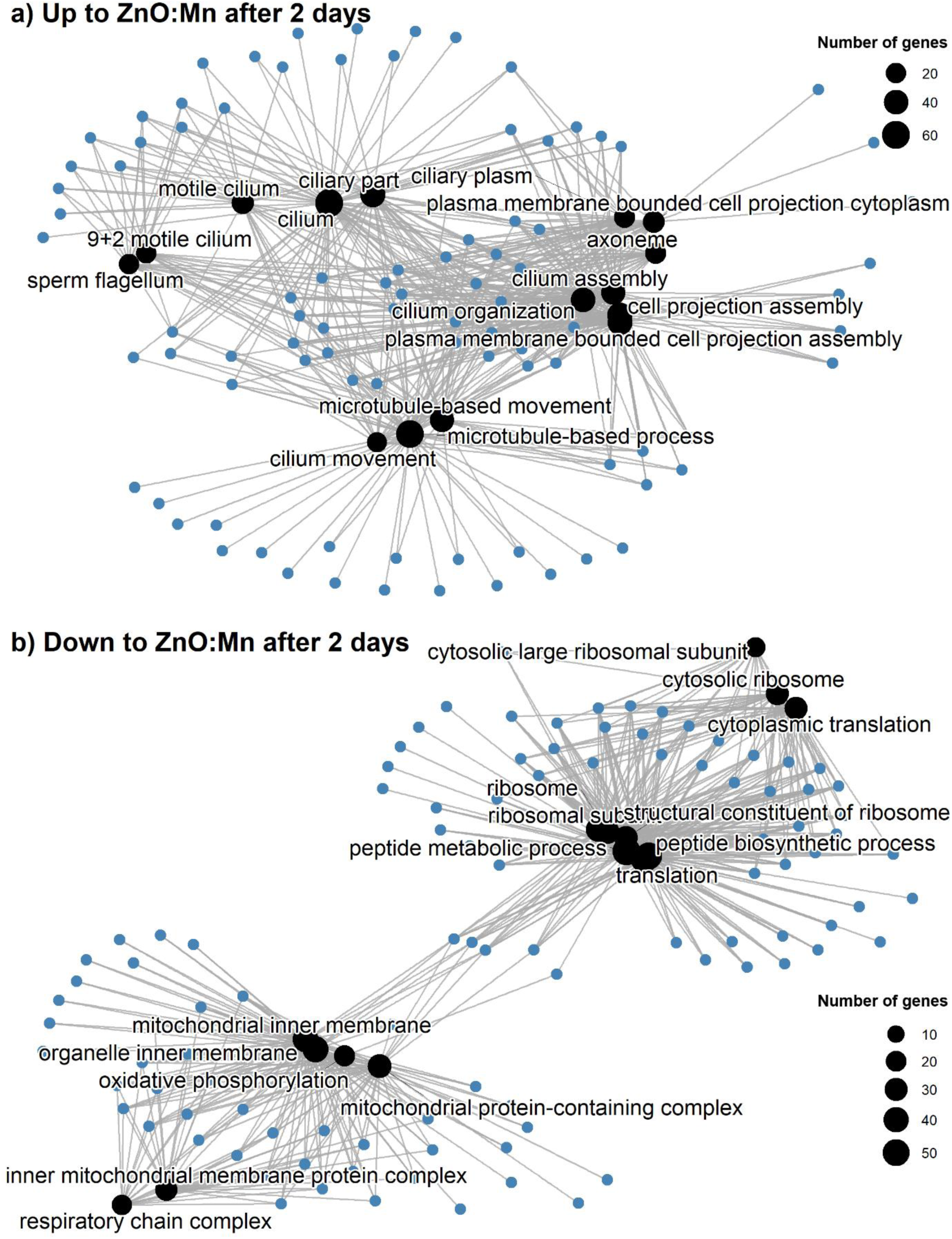
Networks of enriched genes (blue circles) and top 15 gene ontology terms (black circles, size represents the number of genes associated to each enriched GO term) for a) downregulated genes in earthworms exposed to ZnO:Mn for 2 days and b) upregulated genes in earthworms exposed to ZnO:Mn for 2 days in experiment 2. Grey edges connect genes to GO terms.

Finally, upregulated and downregulated genes after 14 days of exposure to ZnO:Mn were enriched in 24 and 50 GO-terms, respectively (File S3). Upregulated and downregulated genes after 14 days of exposure to MnCl2 were enriched in 66 and 5 GO terms, respectively (File S3). Enriched GO terms among the DEG after 14 days of exposure were associated with no more than 4 genes which suggests these are not systemic effects. These results at the endpoint observation after 14 days of exposure to ZnO and ZnO:Mn nanomaterials and MnCl2, may indicate that the applied materials are rather non-toxic to earthworms, which is in contrast to Ag ions or Ag nanomaterials for which toxicity was observed after 28 days of exposure (Novo *et al*., 2015).

## Conclusion

Gene expression profiling in earthworms exposed to ZnO indicated a transient response with pronounced differential gene expression on day 4. Earthworms exposed to ZnO:Mn also indicated a transient response, but with pronounced differential expression on day 2. These temporary peaks in differential gene expression were confirmed by a regression-based time-dependent analysis. Although these transient responses were obtained by mapping reads to *E. andrei* reference genomes, end-point transcriptomic measurements after exposing earthworms to ZnO:Mn and MnCl2 for 14 days partly used a *de novo* assembled transcriptome, which indicated moderate differential gene expression. GO enrichment analysis of genes that were differentially expressed after exposure to ZnO contained terms related to various supramolecular, actin and muscle cell related cellular components. In addition, genes upregulated after 2 days of exposure to ZnO:Mn were associated with cellular components related to cilia, microtubules and cell projection. Both of these two collections of GO terms could be associated with muscle biology. In contrast, genes downregulated after 2 days of exposure to ZnO:Mn were associated with energy metabolism and translation. Obtained GO terms included peptide, oxidative phosphorylation, cytoplasmic translation related biological processes, molecular functions and various cytosolic ribosome, inner mitochondrial and respiratory chain complex cellular components. Finally, GO terms obtained by enriching genes that were differentially expressed after exposing earthworms to ZnO:Mn and MnCl2 for 14 days had only a few genes per GO term. Therefore, exposing earthworms to ZnO NM and ZnO:Mn MCNM elicits a temporary response in differential gene expression that largely disappears by day 14 from the start of the exposure.

## CRediT authorship contribution statement

***H.J. van Lingen***: Conceptualization, Data curation, Formal analysis, Investigation, Methodology, Software, Validation, Visualization, Writing – original draft, Writing – review and editing. ***C. Ke***: Data curation, Formal analysis, Writing – original draft, Methodology. ***E. Saccenti***: Conceptualization, Methodology, Supervision, Validation, Writing – review and editing. ***M. Suarez-Diez***: Conceptualization, Methodology, Funding acquisition, Project administration, Supervision, Validation, Writing – review and editing. ***M. Baccaro***: Investigation, Methodology, Resources, Writing – review and editing. ***N.W. van den Brink***: Conceptualization, Funding acquisition, Project administration, Supervision, Validation, Writing – review and editing. ***Z.G. Lada***: Formal analysis, Investigation, Methodology, Visualization, Writing – original draft

## Acknowledgements

We acknowledge Bart Nijsse (Wageningen University & Research) for his valuable input on the design and optimization of the bioinformatic pipelines employed in this study. In addition, we thank Andreas Stingl and Patricia M.A. Farias (Phornano Holding GmbH, Kleineingersdorfer Straße 24, 2100 Korneuburg, Austria) for providing us the ZnO and ZnO:Mn nanomaterials.

## Declaration of generative AI and AI-assisted technologies in the writing process

During the preparation of this work the author(s) used ChatGPT in order to improve the readability of the writing output generated by the authors. After using this tool, the authors reviewed and edited the content as needed and take full responsibility for the content of the published article.

