## Supplementary Information for "Gene expression profile dynamics of earthworms exposed to ZnO and ZnO:Mn nanomaterials"

### Experimental details of ZnO and ZnO:Mn nanomaterials characterization

Raman spectra were recorded on a T-64000 (Jobin Yvon-Horiba) micro-Raman system equipped with a 2D-CCD Symphony II detector. The excitation wavelength (514.5 nm) was provided by a DPSS laser (Cobolt Fandango TMISO laser). Dispersion and detection of the Raman photons were done by a 600 grooves/mm grating and a Spectraview-2DTM liquid N2-cooled CCD detector, respectively. The laser was focused on the samples by a 50x microscope objective with a power of 1 mW on sample. The resolution was kept constant in all experiments (~4 $cm^{-1}$). XRD spectra were performed for the structural characterization of ZnO and ZnO:Mn samples by using a Bruker D8 Advance diffractometer equipped by a Cu lamp (λCuKa = 1.54046 Å) at a scanning rate 0.5 sec/step over a range 2–80$^{o}$ (2θ). SEM images were obtained using a Zeiss SUPRA 35VP-FEG instrument, operating at 5–20 keV. The system is equipped with EDX (Bruker GmbH, Quanta 200) and BSE (K E Developments), which enabled elemental analyses.

### Workflow for processing of reads and quality control for all samples

Docker containers were used throughout to ensure reproducibility. Initial and after-filtering quality check was performed using FastQC v0.11.9 (Andrews, 2010). Reads were then trimmed with Fastp v0.20.1 (Chen et al., 2018), which corrected mismatched base pairs in overlapped regions of paired-end reads to improve overall read accuracy. Additionally, reads shorter than a 50 bp were removed. Low-quality bases were filtered out by requiring minimum Phred score Q20 and reads with more than 20% low-quality bases were discarded. BBduk v38.87 (Bushnell, 2014) was used to remove non-target sequences such as ribosomal RNA (rRNA) and phiX. To check for potential contaminants, taxonomic classification was performed using Kraken2 with full PlusPFP-16 database, followed by refinement of abundance estimates using Bracken. The results from these quality controls and all subsequent analyses were aggregated using multiQC v1.9 (Ewels et al., 2016).

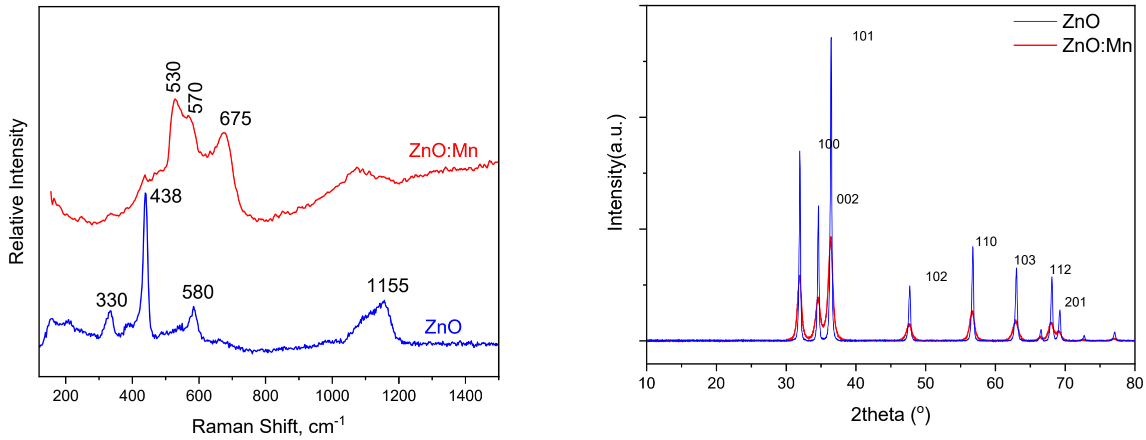


Figure S1. Raman spectra (a) and X-ray diffraction (b) of ZnO and ZnO:Mn samples.

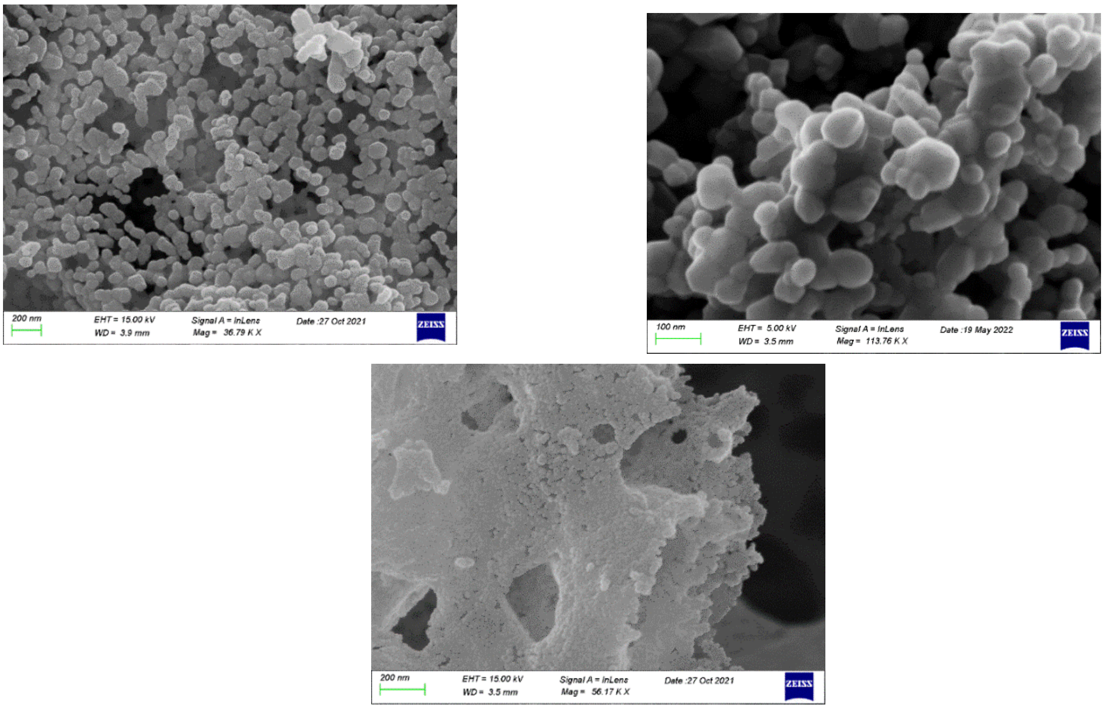


Figure S2. Representative SEM images of samples.

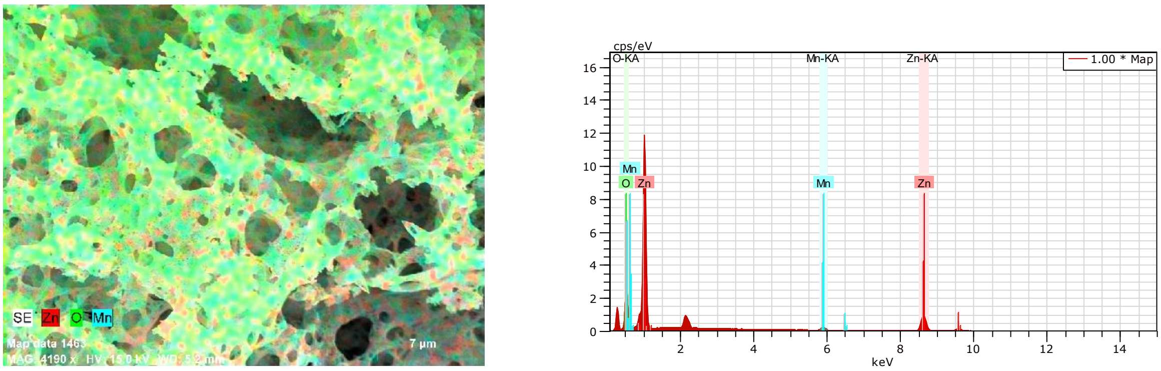


Figure S3. SEM-EDX images for ZnO:Mn sample.

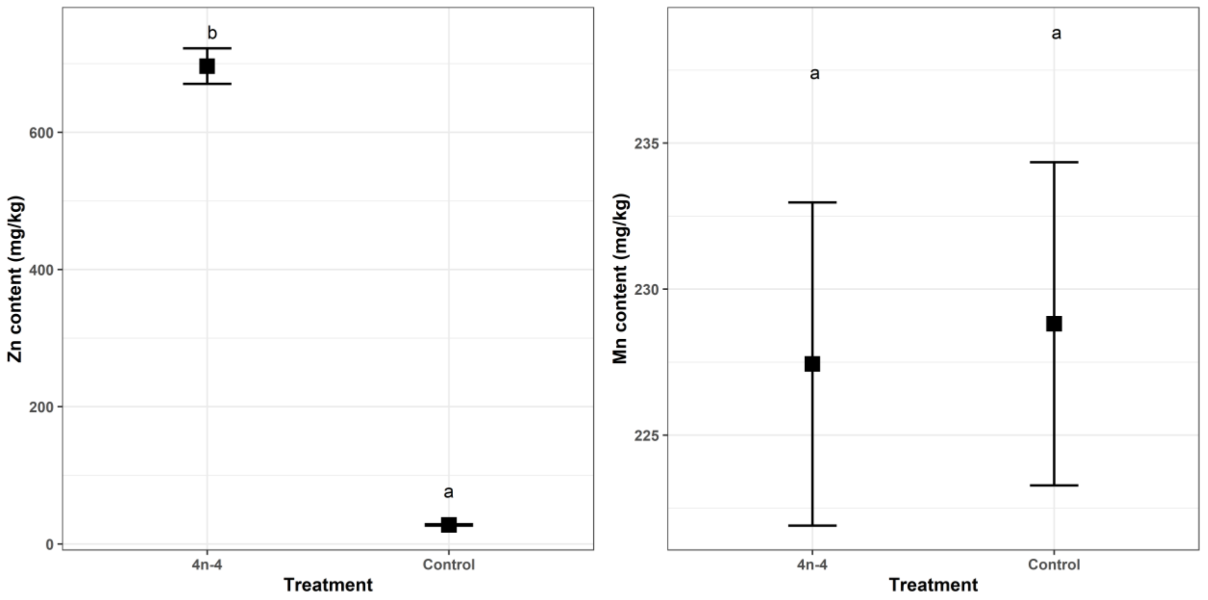


Figure S4 – Zinc and manganese contents (mg/kg) in samples of ZnO treated and control soil in experiment 1.

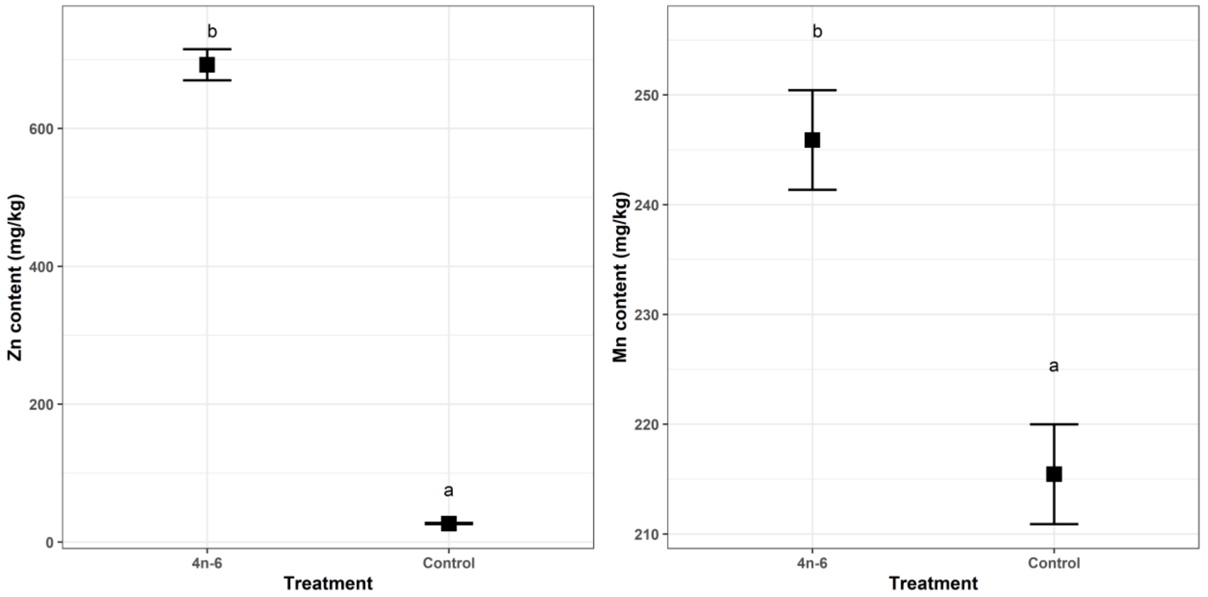


Figure S5 – Zinc and manganese contents (mg/kg) in samples of ZnO:Mn treated and control soil samples in experiment 2.

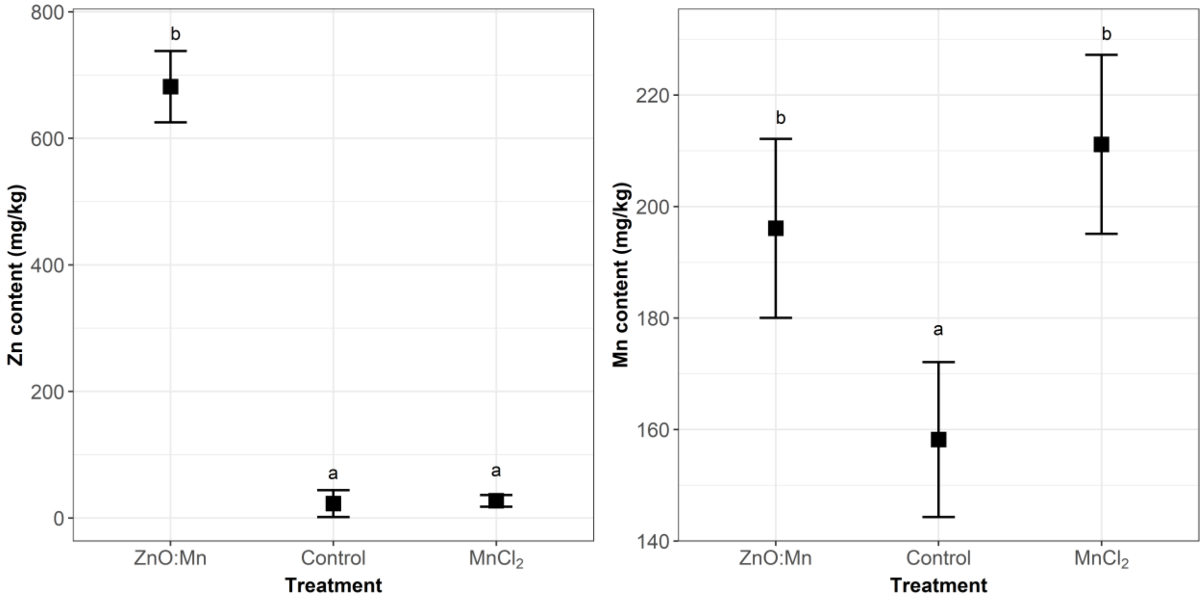


Figure S6 – Zinc and manganese contents (mg/kg) in samples of ZnO:Mn treated, control treated and $MnCl_2$ treated soil samples in experiment 3.

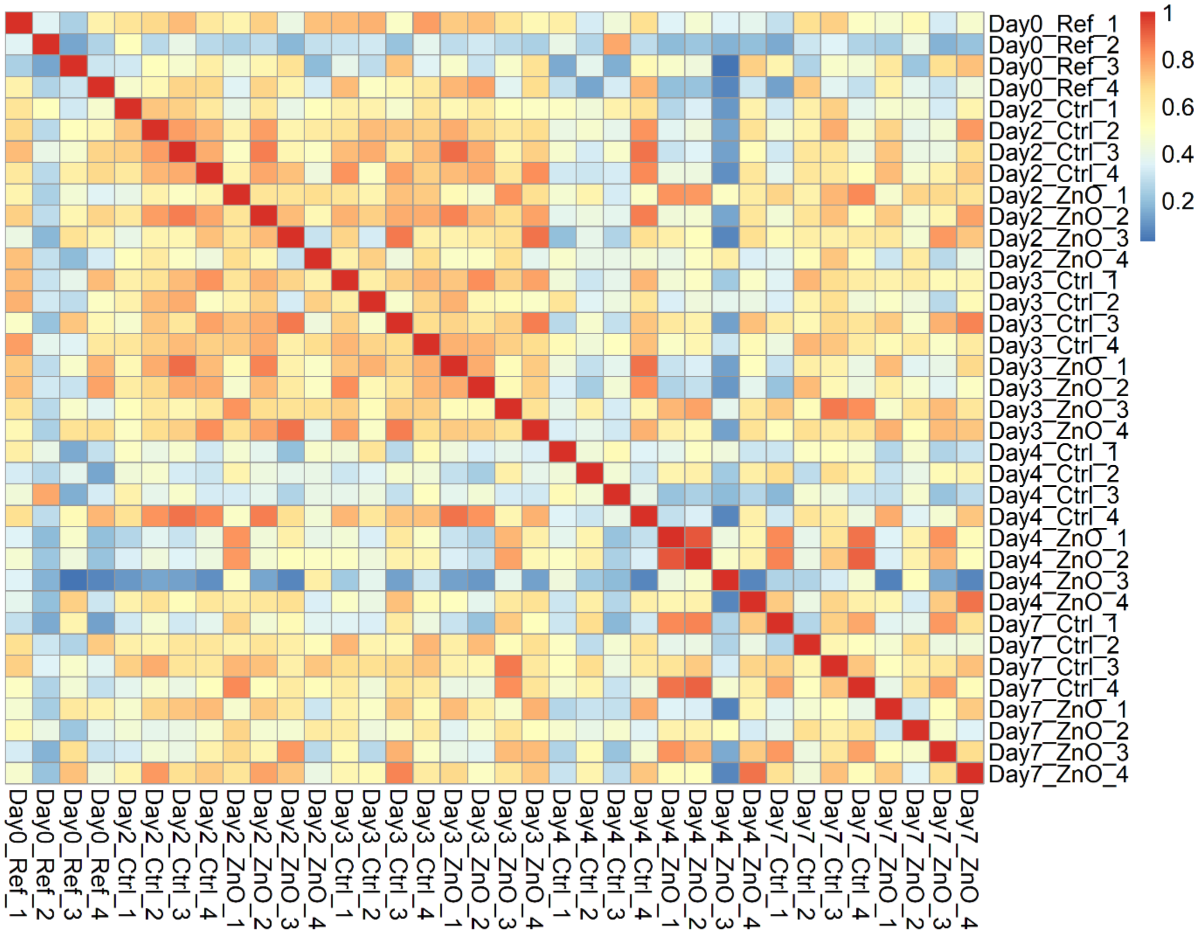


Figure S7 – Heatmap for correlations of ~29,000 gene transcripts mapped to *E. andrei* for experiment 1.

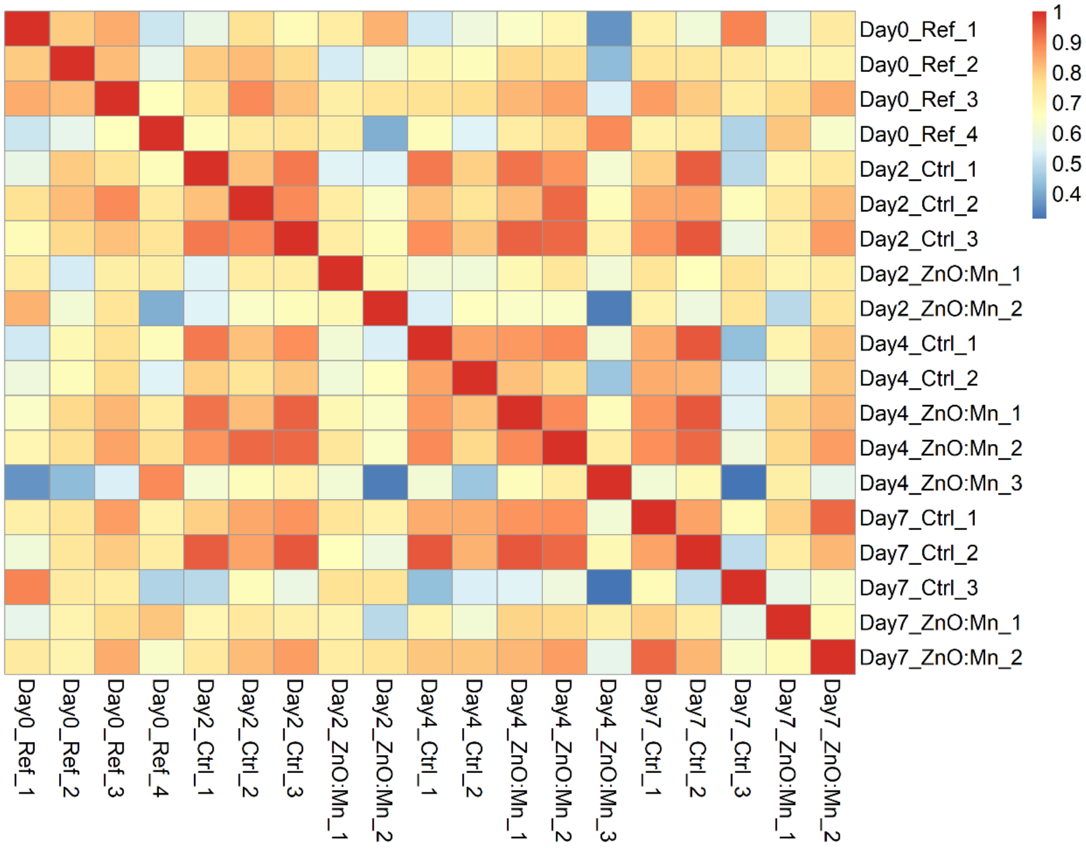


Figure S8 – Heatmap for correlations of ~29,000 gene transcripts mapped to *E. andrei* for experiment 2.

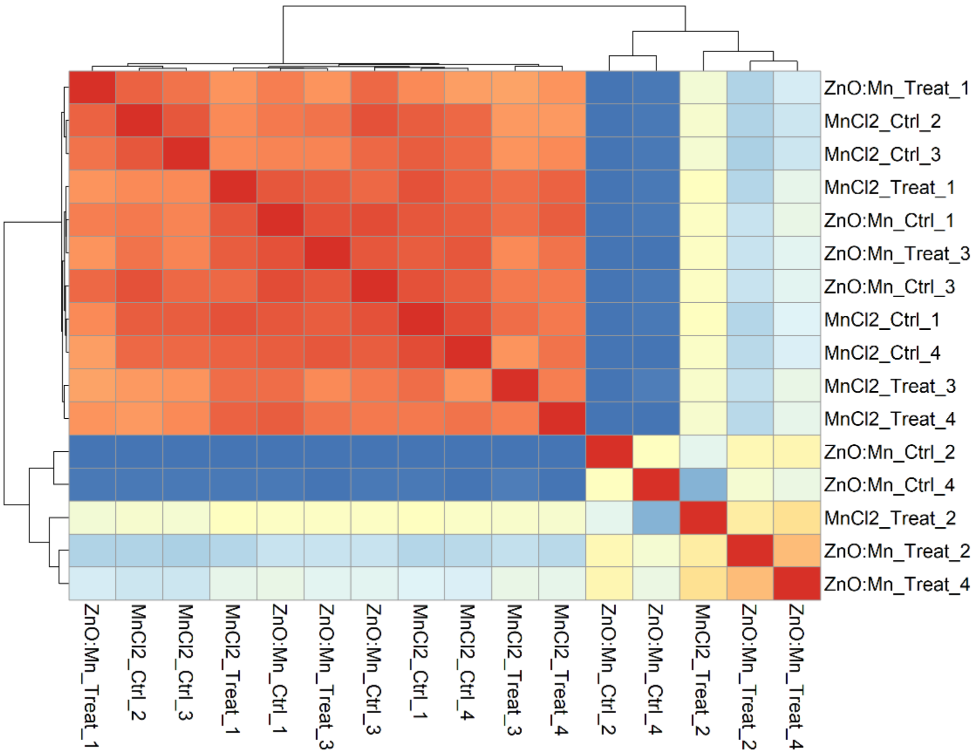


Figure S9 – Heatmap for correlations of ~73,000 *de novo* assembled gene transcripts for experiment 3.

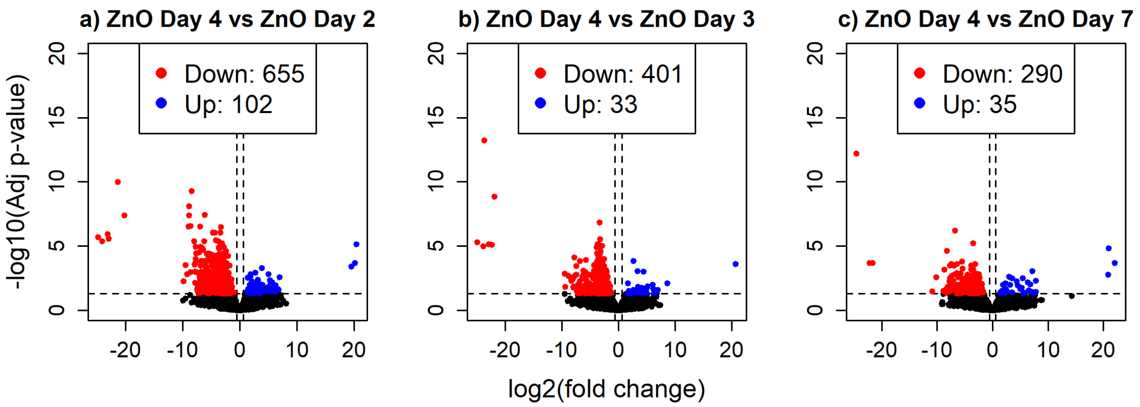


Figure S10 - Volcano plots that represent differential gene expression of transcripts mapped to the *E. andrei* reference genome for earthworms exposed to ZnO treated soil for 4 days *vs.* earthworms exposed to ZnO treated soil for 2, 3 and 7 days in experiment 1 (a-c).
